# Transcriptome analysis of developmental stages of cocoa pod borer, *Conopomorpha cramerella*: A polyphagous insect pest of economic importance in Southeast Asia

**DOI:** 10.1101/2021.06.01.446533

**Authors:** Chia Lock Tan, Rosmin Kasran, Wei Wei Lee, Wai Mun Leong

## Abstract

The cocoa pod borer, *Conopomorpha cramerella* (Snellen) is a serious pest in cocoa plantations in Southeast Asia. It causes significant losses in the crop. Unfortunately, genetic resources for this insect is extremely scarce. To improve these resources, we sequenced the transcriptome of *C. cramerella* representing the three stages of development, larva, pupa and adult moth using Illumina NovaSeq6000. Transcriptome assembly was performed by Trinity for all the samples. A total number of 147,356,088 high quality reads were obtained. Of these, 285,882 contigs were assembled. The mean contig size was 374 bp. Protein coding sequence (CDS) was extracted from the reconstructed transcripts by TransDecoder. Subsequently, BlastX and InterProScan were applied for homology search to make a prediction of the function of CDS in unigene. Additionally, we identified a number of genes that are involved in reproduction and development such as genes involved in general function processes in the insect. Genes found to be involved in reproduction such as *porin, dsx, bol* and *fruitless* were associated with sex determination, spermatogenesis and pheromone binding. Furthermore, transcriptome changes during development were analysed. There were 2,843 differentially expressed genes (DEG) detected between the larva and pupa samples. A total of 2,861 DEG were detected between adult and larva stage whereas between adult and pupa stage, 1,953 DEG were found. In conclusion, the transcriptomes could be a valuable genetic resource for identification of genes in *C. cramerella* and the study will provide putative targets for RNAi pest control.

## Introduction

Cocoa pod borer (*Conopomorpha cramerella* Snellen) is a Lepidopteran moth of the family *Gracillariidae* [1]. It is known to be of south Asian origin [2]. It is found mainly in Thailand, Brunei, Indonesia (Sumatra, Sulawesi, Papua New Guinea, Java, Kalimantan, Moluccas), Malaysia, Vietnam, Australia, Philippines, Samoa, the Solomon Islands, Sri Lanka, Taiwan and Vanuatu. Its primary hosts are plants native to the area such as Rambutan (*Nephelium lappaceum*); Pulasan (*Nephelium mutabile*); Kasai (*Pometia pinnata*); Cola (*Cola nitida*, *C. acuminate*); and Nam-nam, (*Cynometra cauliflora*). With the introduction of cocoa (*Theobroma cacao* L.) to this geographic region, cocoa pod borer (CPB) moved onto this crop and exploited *T. cacao* as its new host. Since 1986, CPB has become the most serious insect pest of cocoa in Southeast Asia (Indonesia, Philippines, Malaysia, and Papua New Guinea). Economic losses due to this insect can be up to 80% in some geographical regions [56]. Control of this notorious pest is achieved mainly by chemical pesticides. However, overuse of pesticides leads to environmental and food safety issues. Therefore, alternate pest control strategies for CPB is highly desirable and need to be developed.

Double-stranded RNA (dsRNA)-mediated gene silencing, commonly referred to as RNA interference (RNAi), is becoming a widely used functional genomics tool in insects to ascertain the function of the many newly identified genes accumulating from genome sequencing projects [3, 4]. The basic components of the RNAi process, namely the endonuclease Dicer, which first chops long dsRNAs into short interfering RNAs (siRNAs), and the RNA-induced silencing complex (RISC), which facilitates the targeting and endonucleolytic attack on mRNAs with sequence identity to the dsRNA, are evolutionarily conserved across virtually all eukaryotic taxa [5], and consequently, RNAi could be readily applied to any insect species. This RNAi technique has been successfully applied to study gene functions in many insects, including *Drosophila melanogaster* [6], *Tribolium castaneum* [7], *Helicoverpa armigera* [8], *Gryllus bimaculatus* [9], *Schistocerca gregaria* [10], *Plutella xylostella* [11], *Nilaparvata lugens* [12], and *Epiphyas postvittana* [13]. There are two kinds of RNA delivery methods, oral intake or injection. Injection of siRNA or dsRNA is widely used in the laboratory at a small scale level, whereas oral intake is more feasible to be used for controlling pest in the field condition.

RNAi-mediated pest control is a novel and promising technique because interference with important insect genes using RNAi can lead to death of pests. Proof of principle for the application of RNAi in insect crop pest control comes from early studies conducted on the western corn rootworm (WCRW - *Diabrotica virgifera*), and cotton bollworm (CBW - *Helicoverpa armigera*) [14]. The researchers fed larval WCRW on 290 dsRNAs, from which they identified 14 genes that reduced larval performance, and one of these, vacuolar ATPase subunit A (V-ATPase), was carried forward for detailed analysis. Low concentrations of orally-delivered dsRNA against V-ATPase in artificial diet suppressed the corresponding WCRW mRNA. Importantly, larvae reared on transformed corn plants that express V-ATPase dsRNA also displayed reduced expression of the V-ATPase gene and caused much reduced plant root damage. In the study on CBW, the target gene was a cytochrome P450, CYP6AE14, which is expressed in the larval midgut and detoxifies gossypol, a secondary metabolite common to cotton plants. When CBW was exposed to either *Arabidopsis thaliana* or *Nicotiana tobacum* expressing CYP6AE14 dsRNA, levels of this transcript in the insect midgut decreased, larval growth was retarded, and both effects were more dramatic in the presence of gossypol. Transgenic cotton plants expressing CYP6AE14 dsRNA also support drastically retarded growth of the CBW larvae, and suffered less CBW damage than control plants [15]. In another study, researchers used hairpin RNA expressed in both *Escherichia coli* and transgenic tobacco plants to decrease mRNA and protein levels of the *H. armigera*- derived molt-regulating transcription factor in larval *H. armigera*, which resulted in developmental deformity and larval lethality [16]. Another example is provided by nicotine, a neurotoxin made by species of tobacco. The tobacco hornworm *Manduca sexta* (Lepidoptera) can tolerate high nicotine concentrations. Larvae even exhale nicotine through their spiracles, deterring spider predation. Dietary nicotine induces the cytochrome P450 gene CYP6B46 in *M. sexta*. Tobacco plant transformed with a construct expressing dsRNA targeting 300 nt of the *M. sexta* gene for CYP6B46. Tobacco hornworm larvae consuming the transformed tobacco were more susceptible to spider predation because they exhaled less nicotine [17]. The success of these studies attests to the functionality of the RNAi in controlling insect pests.

To develop RNAi-mediated pest control methods, it is critical to find suitable target genes. Target genes should not only have insecticidal effects on the target pests, but should also be safe to non-target organisms. Unfortunately, genetic resources for CPB insect is extremely scarce and therefore additional resources are required for effective screening of target genes. Insect transcriptomes has been reported to be useful genetic resources for high-throughput screening of RNAi target genes [18]. The introduction of next-generation sequencing technologies has provided significant convenience for further studies of non-model organisms including insects [19, 20]. Next generation sequencing such as Illumina and PacBio have been widely used to identify genes involved in several developmental and physiological processes. These technologies has been used to identify candidate chemosensory genes of oligophagous insect, *Ophraella communa* (Coleoptera: Chrysomelidae). These genes plays a key role in insect survival, which mediates important behaviors like host search, mate choice, and oviposition site selection [21]. Using NextSeq500 (Illumina) sequencing, Singh *et al*. [22] studied *de novo* transcriptome assembly and analysis of RNAi in *Phenacoccus solenopsis* Tinsley (Hemiptera: Pseudococcidae), one of the major polyphagous crop pests in India. The study provides a base for future research on developing RNAi as a strategy for management of this pest. Gao *et al.* [23] used PacBio to profile full-length transcriptomes of insect *Erthesina fullo* Thunberg mitochondrial gene expression. However, even though CPB is an important pest to cocoa in South-east Asia, there is no published report on the genome or transcriptome of the insect. To the best of our knowledge, this is the first report on transcriptomic analysis of *C. cramerella*, covering the three developmental stages of the life cycle of the insect.

In this study, we present the results from the sequencing and assembly of the transcriptome of *Conopomorpha cramerella* Snellen at different developmental stages (larvae to pupa and adult) using Illumina NovaSeq6000 technology. Genes involved in metabolic processes, general development and reproduction were identified and functionally annotated. A great number of differentially expressed genes were obtained and some of these genes have been cloned using PCR for further downstream studies. The transcriptome study is undoubtedly valuable for molecular studies of the underlying mechanism on the development and reproduction of the insect. It also serve as a useful resource for target genes for RNA interference studies and the development of effective and environmental-friendly strategies for pest control.

## Materials and methods

### Insects

Cocoa pods that were infected with cocoa pod borer (CPB) were obtained from cocoa farm in Keningau, Sabah, Malaysia. They were wrapped in papers and kept in the dark for two weeks. During the period, they were constantly checked for CPB larvae, pupae and moth.

Approximately thirty of each larvae, pupae and moth were collected and kept in RNA Later® and maintained in -70oC freezer until later use. The samples were grind to fine powder in liquid nitrogen before RNA isolation.

### RNA isolation and cDNA construction

Total RNA from CPB larvae, pupae and moth were extracted using the GeneAll Hybrid-R™ kit (GeneAll Biotechnology, Seoul, Korea) according to the manufacturer’s instructions. RNA Integrity Number (RIN) was determined using RNA Nano 6000 Assay Kit (Agilent Technologies, CA, USA) with the Agilent 2100 Bioanalyzer (Agilent Technologies).

The libraries were prepared for 150bp paired-end sequencing using TruSeq stranded mRNA Sample Preparation Kit (Illumina, CA, USA). Namely, mRNA molecules were purified and fragmented from 1μg of total RNA using oligo (dT) magnetic beads. The fragmented mRNAs were synthesized as single-stranded cDNAs through random hexamer priming. By applying this as a template for second strand synthesis, double-stranded cDNA was prepared. After sequential process of end repair, A-tailing and adapter ligation, cDNA libraries were amplified with PCR (Polymerase Chain Reaction). Quality of these cDNA libraries was evaluated with the Agilent 2100 BioAnalyzer (Agilent, CA, USA). They were quantified with the KAPA library quantification kit (Kapa Biosystems, MA, USA) according to the manufacturer’s library quantification protocol. Following cluster amplification of denatured templates, sequencing was progressed as paired-end (2×150bp) using Illumina NovaSeq6000 (Illumina, CA, USA).

### Bioinformatics Analysis of RNA-seq data: Transcriptome assembly & Unigene discovery

#### A. Filtering

Prior to the assembly, filtering was proceeded to remove low quality reads and adapter sequence according to the following criteria; reads contain more than 10% of skipped bases (marked as ‘N’s), reads contain more than 40% of bases whose quality scores are less than 20 and reads of which average quality scores of each read is less than 20. Furthermore, bases of both ends less than Q20 of filtered reads were removed additionally. This process is to enhance the quality of reads due to mRNA degradation in both ends of it as time goes on [24]. The whole filtering process was performed using the in-house scripts.

#### B. Assembly

Transcriptome assembly was performed by Trinity [25, 26] program using data from all samples. Trinity is a representative RNA assembler based on the de Bruijin graph (DBG) algorithm for RNA-seq de novo assembly, and its assembly pipeline consists of three consecutive modules: Inchworm, Chrysalis, and Butterfly. First, Inchworm module is to construct contigs according to the following steps; each 100bp read divides into 4 fragments (each fragment is 25bp). When to overlap 24bp of the each fragment, the 24 overlapped region is merged for construction of contigs. The module requires a single high-memory server so that classification into subgroups after the construction was progressed for efficient usage of memory. Next, Chrysalis clusters related Inchworm contigs into components. And, the DBG is generated in each cluster. Finally, Butterfly reconstructs transcript sequences in a manner that indicates the original cDNA molecules. All options were set to default values.

#### C. Clustering

According to the previous publication [27], there are some problems as to when to perform the assembly by Trinity. At first, the assembled transcripts contained the overlapping sequence of same region. This is due to the transcripts originated from transcripts containing isoforms and not genes. In addition to that, chimera transcripts are generated through the assembly process. To overcome these problems, grouping the assembled transcripts by TGICL [28], a pipeline for transcriptome analysis in which the sequences are clustered based on pairwise sequence similarity, was carried out for removal of the overlapping and the chimera sequences. Subsequently, extraction of the representative sequence was carried out using CAP3 [29]: a sequence assembly program. The criterion of sequence similarity for grouping was set to 0.94 value.

#### D. CDS prediction

Protein coding sequence (CDS) was extracted from the reconstructed transcripts by TransDecoder: a utility included with Trinity to assist in the identification of potential coding regions [26]. The coding region is predicted according to following procedures; 1) search all possible CDSs of the transcripts, 2) verify the predicted CDSs by GeneID [30] through selecting it for more than 0 value of log-likelihood score, and 3) choose the region which has the highest score among candidate sequences.

### Functional annotation of Unigenes

Blast and InterProScan were applied for homology search to make a prediction of the function of CDS in unigene.

#### A. Blastx with nucleotide sequence

NCBI Blast 2.2.29+ was applied for nucleotide sequence-based homology search. The function of CDS was predicted by Blastx to search all possible proteins matched with unigene sequence against the SwissProt db. The criterion regarding significance of the similarity was set to E-value < 1e-5.

#### B. InterProScan with protein sequence

InterProScan is another tool for homology search using protein sequence. The InterProScan is based on Hidden Markov Model to predict the function of CDS by similarity search using the protein domain: units of protein structure for function. The search was progressed by InterProScan v5 against ProDom, PfamA, Panther, SMART, SuperFamily and Gene3d databases based on E-value < 1e-5.

### Gene expression estimation

Gene expression level was measured with RSEM [31]. The RSEM is a tool to measure the expression for transcripts without any information on reference, and Bowtie is applied to the RSEM using directed graph model following reads alignment to the transcripts for the expression.

### Differential Expressed gene (DEG) analysis

TCC package was applied for DEG analysis through the interative DEGES/DEseq method. This method is based on DESeq [32] using Negative-binomial distribution. Normalization was progressed three times to search meaningful DEGs between comparable samples [33].

The DEGs were identified based on the qvalue threshold less than 0.05.

### Data availability

The datasets generated and analysed during the current study are available at NCBI Gene Expression Omnibus (GEO) Accession Series GSE146610.

### qRT-PCR validation

To verify the differential expression detected by Illumina RNA-Seq, qRT-PCR was performed on the same samples that had been used previously. A set of seventeen genes was chosen at random, the expression of each gene was evaluated for two life stages of cocoa pod borer insect and compared with their observed FPKM. qRT-PCR was performed using Rotor- Gene™ 6000 Real-Time thermocycler (Corbett Research, Australia) with Brilliant SYBR® Green QPCR Master Mix (Stratagene, La Jolla, CA) following the manufacturer’s instructions. The forward and reverse primers used for qRT-PCR are listed in Supplementary Table S6. The thermal cycling conditions were as follows: 95oC for 10 min, followed by 45 cycles of 95oC for 15 s and 55oC for 60oC. Gene expression was normalised with actin gene using primer pairs qActin-F and qActin-R (Supplementary Table S6). The data are presented as mean ± SE of three independently produced RT preparations used for PCR runs, each having at least three replicates. The relative expression levels were calculated using the delta-delta Ct method [34].

## Results and Discussion

### Generation and assembly of cocoa pod borer transcriptomes

In order to obtain an overview of *Conopomorpha cramerella* gene expression profile, cDNA from three different developmental stages (larvae, pupae and adult moth) were prepared and sequenced on Illumina NovaSeq6000 machine. A total of 22,961,926,438 bp from 147,356,088 sequence reads with an average read length of 146 bp was obtained (Table 1). These raw data were assembled into 285,882 contigs. The mean contig length is 374 bp with lengths ranging from 225 bp to 16,526 bp. The percentage and number of singletons for larvae were 2.88% (1,659,115), pupae: 2.55% (1,044,176) and moth: 2.69% (1,314,145). The GC percentage of the transcriptomes is 38%.

**Table 1.** Summary of the *C. cramerella* transcriptome

|  |  |
| --- | --- |
| Total base pair (bp) | 22,961,926,438 bp |
| Number of high-quality reads | 147,356,088 |
| Number of reads assembled in contigs | 147,356,088 |
| Average read length (bp): | 146 bp |
| Number of contigs | 285,882 |
| Average contig length (bp) | 374 bp |
| Range of contig length (bp): | 225~16,526bp |
| Number of singletons (based on mapped reads counts on the assembled unigene by using BWA software) |  |
| o LARVA: | mapping 72%, singletons 2.88%<br>(1,659,115) |
| o PUPAE | mapping 69.44%, singletons 2.55%<br>(1,044,176) |
| o MOTH | mapping 70.54%, singletons 2.69%<br>(1,314,145) |
| GC percentage | 38% |

### Annotation of predicted sequences

To analyse which part of the assembled sequences had counterparts with other insect species, orthologous genes shared between *C. cramerella* and other three insect species were compared. These insect species chosen for comparison were Dipteran *Drosophila melogaster,* Lepidopteran *Bombyx mori* and Lepidopteran *Helicoverpa armigera.* The results showed a total number of 16,595 hits (Figure 2). There were 7,523 identifiable genes shared between *D. melanogaster* and *C. cramerella*, indicating a good coverage of *C. cramerella* transcriptomes. Homologous genes shared between *Bombyx mori* and *C. cramerella* were 6,047 and between *Helicoverpa armigera* and *C. cramerella* were 5,036. There were more genes that *C. cramerella* shared with both *Bombyx mori* and *Helicoverpa armigera* (1,845) than *C. cramerella* with all the three insects combined (73). This is unsurprising as *C. cramerella, Bombyx mori* and *Helicoverpa armigera* are Lepidopteran insects whereas *Drosophila melogaster* belongs to Diptera.

**Figure 1.**
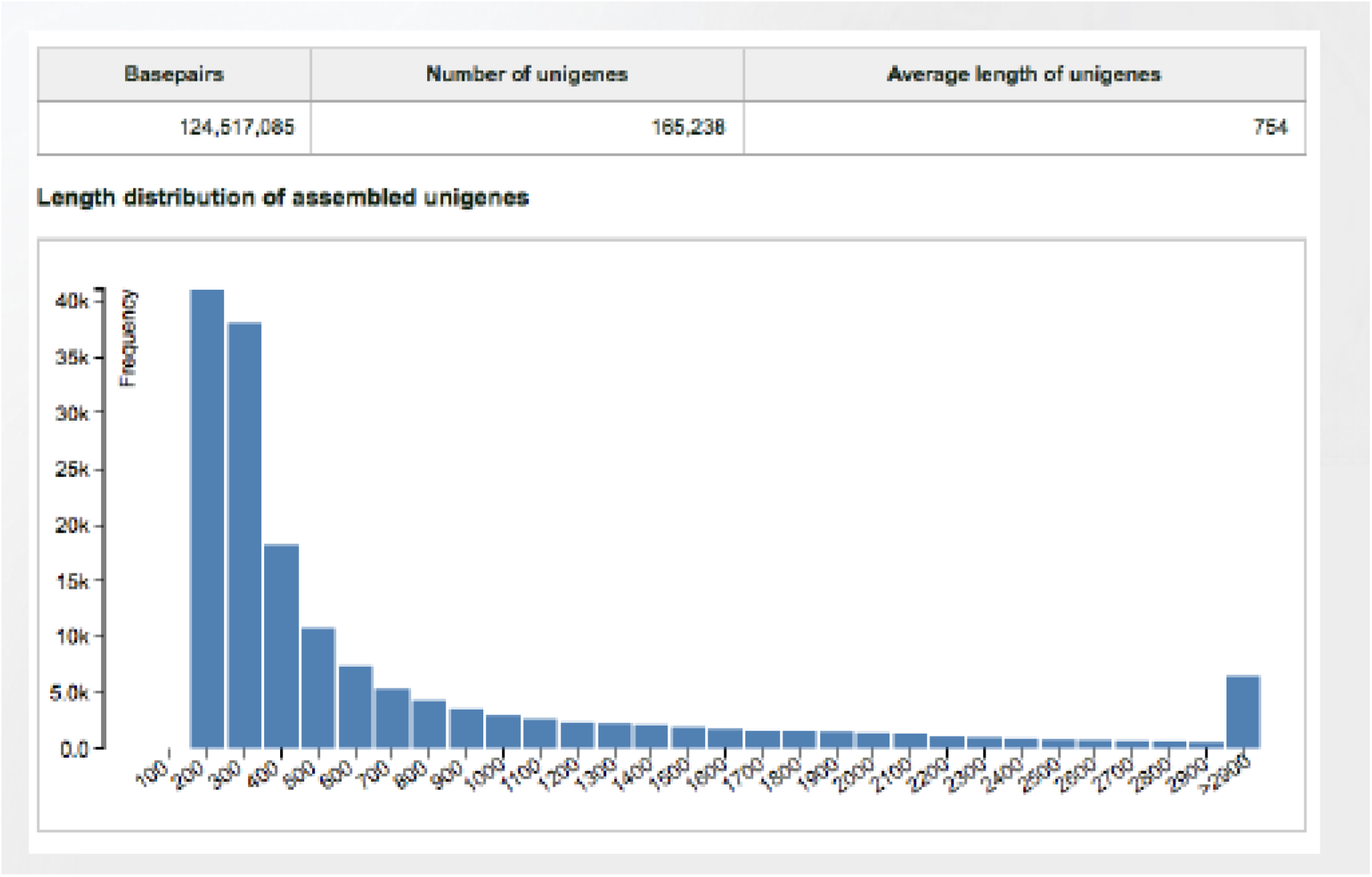
Assembled unigene length distribution of *C. cramerella* transcriptome. The x-axis indicates unigene size and the y-axis indicates the number of unigenes of each size.

**Figure 2.**
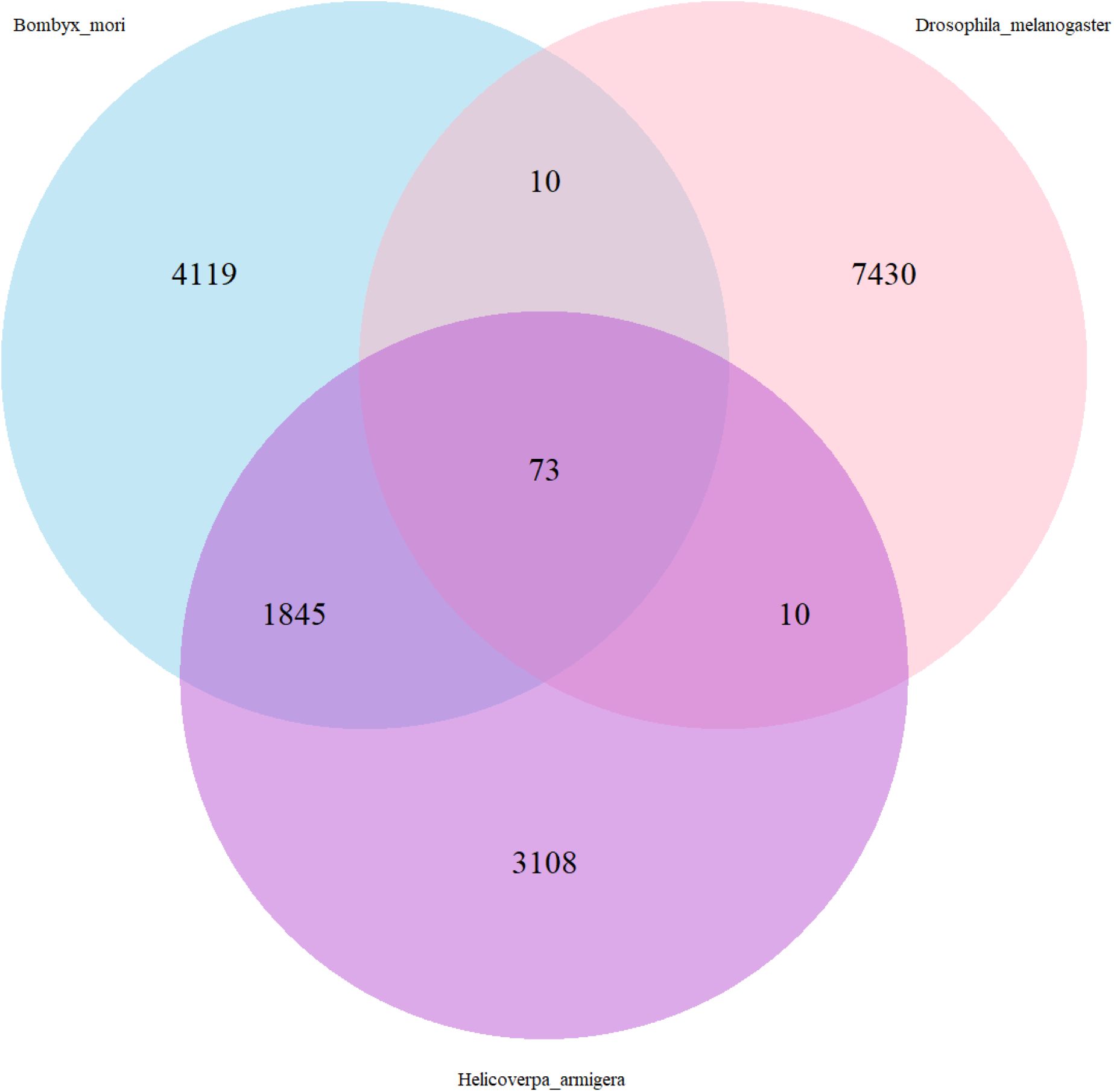
Orthologous gene groups shared between *Conopomorpha, Drosophila, Bombyx* and *Helicoverpa*. Venn diagram of the distribution of the orthologous gene groups among the mentioned species.

The identity distribution of *C. cramerella* transcriptomes were then analysed (Figure 3). Out of a total of 67,770 (41%) hits that has homology, 80.32% (54,434) were of plant origin. The second largest group were invertebrates, which include insects (11.16%, 7,565). The other groups like bacteria, primates, virus and vertebrates were less than 5% of homology.

**Figure 3.**
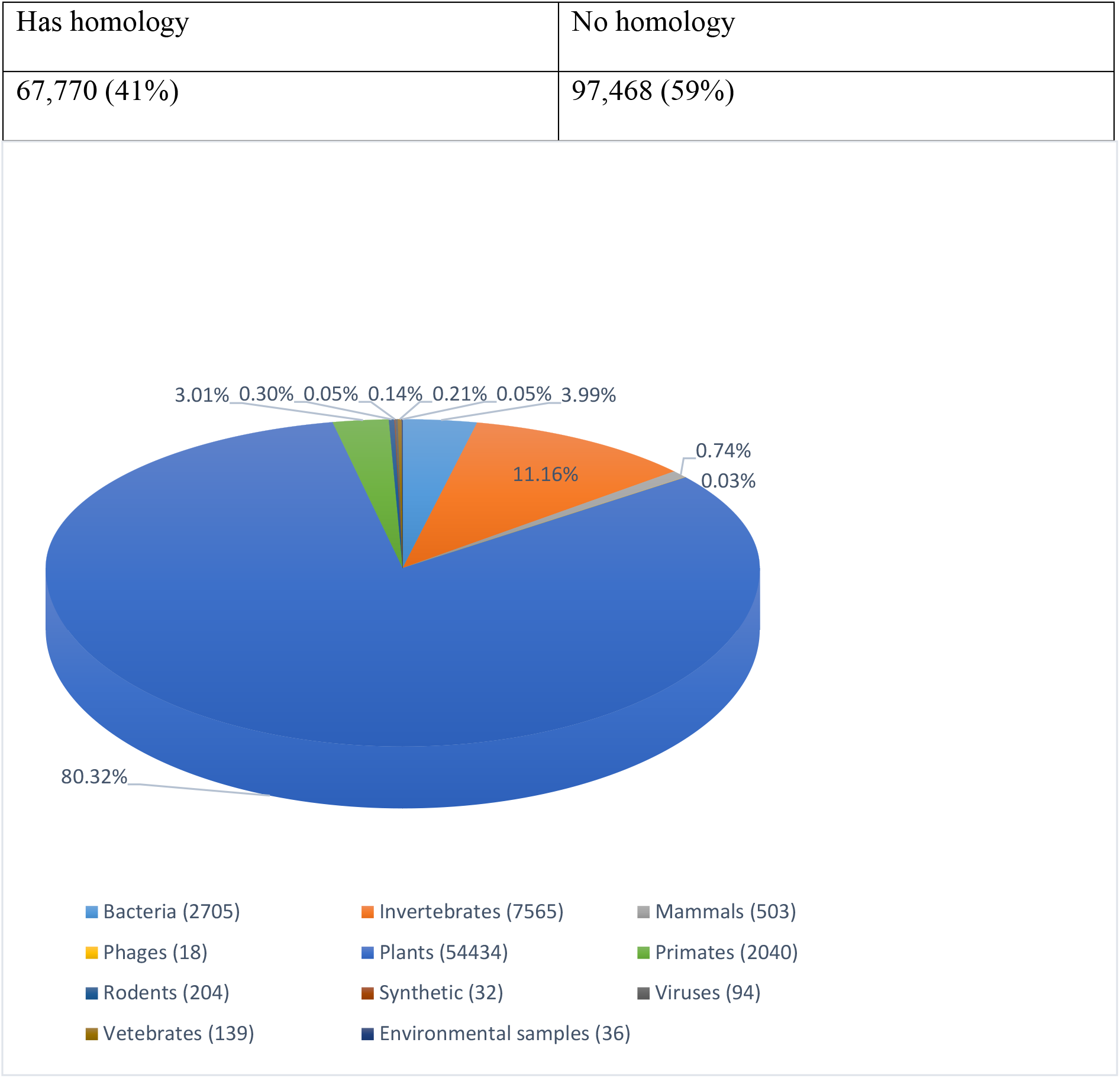
Identity distribution of the top BLAST hits for each sequence of total 67,770 that has homology.

The high homology with plant genes could be due to the fact that *C. cramerella* is a phytophagous insect [35].

### Gene ontology and cluster of orthologous groups classification

Gene ontology (GO) assignment programs were utilised for functional categorisation of annotated genes. These sequences were categorised into 54 main functional groups belonging to 3 categories, including biological process, molecular function and cellular component. Among the biological processes (Figure 4A), the dominant GO terms were grouped into either metabolic process (28%), biological regulation (18%) or cellular process (16%) (Figure 3). Within the molecular function category, there was a high percentage of genes with binding (45%) and catalytic activity (35%) (Figure 4B). For cellular components, those assignments were mostly given to cell part (27%), organelle (21%), membrane part (14%) and membrane (12%) (Figure 4C). The three largest functional groups were binding, catalytic activity and metabolic process.

**Figure 4.**
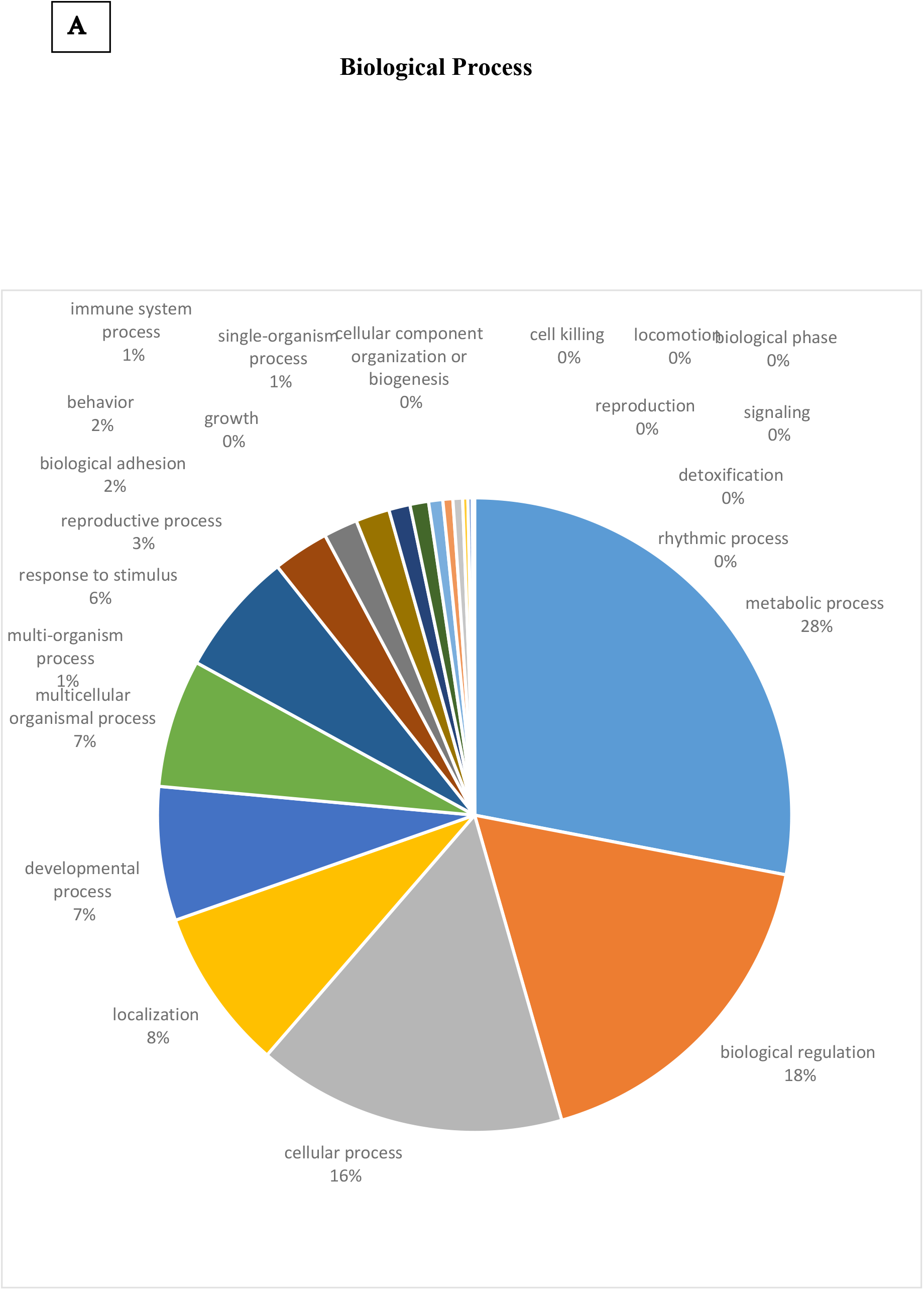
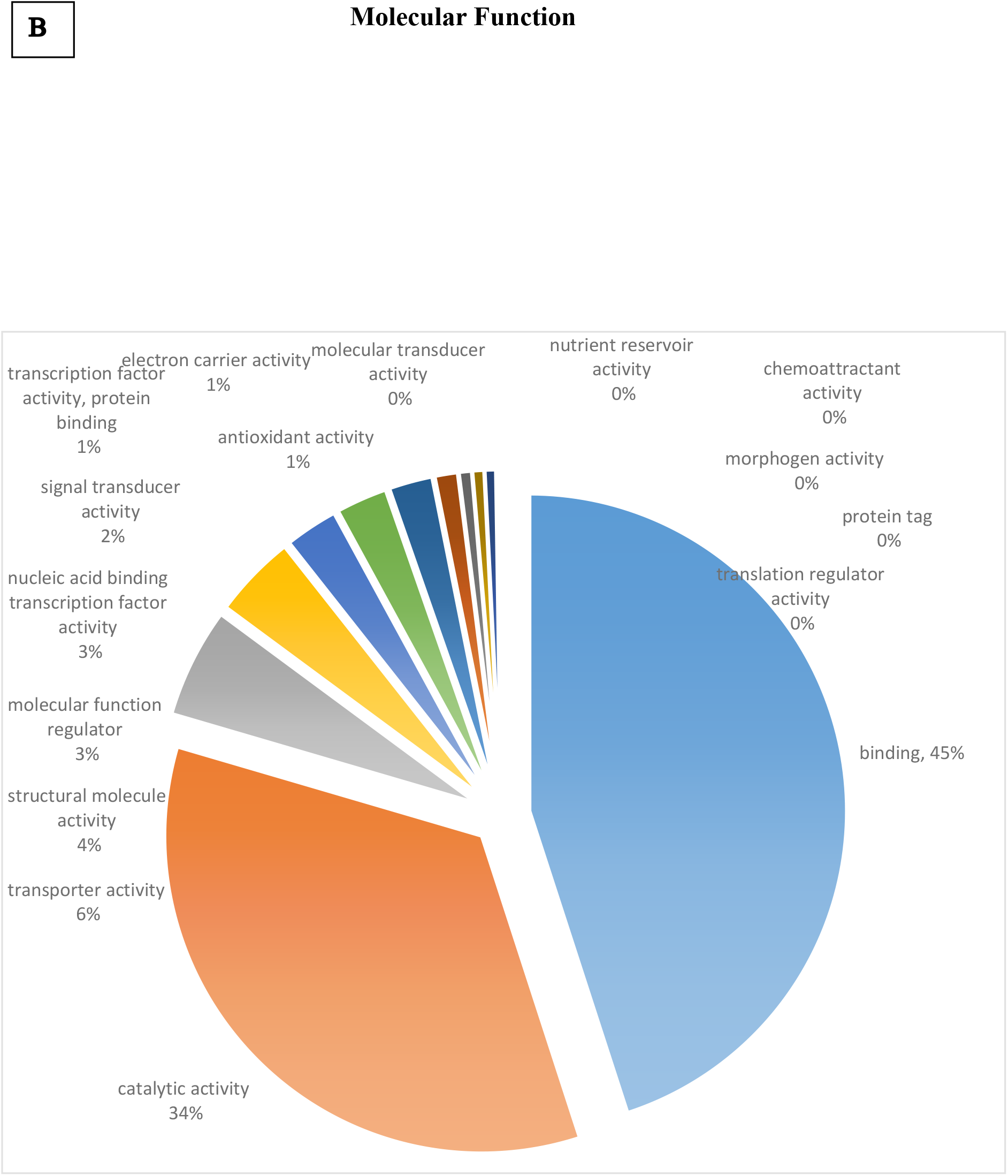
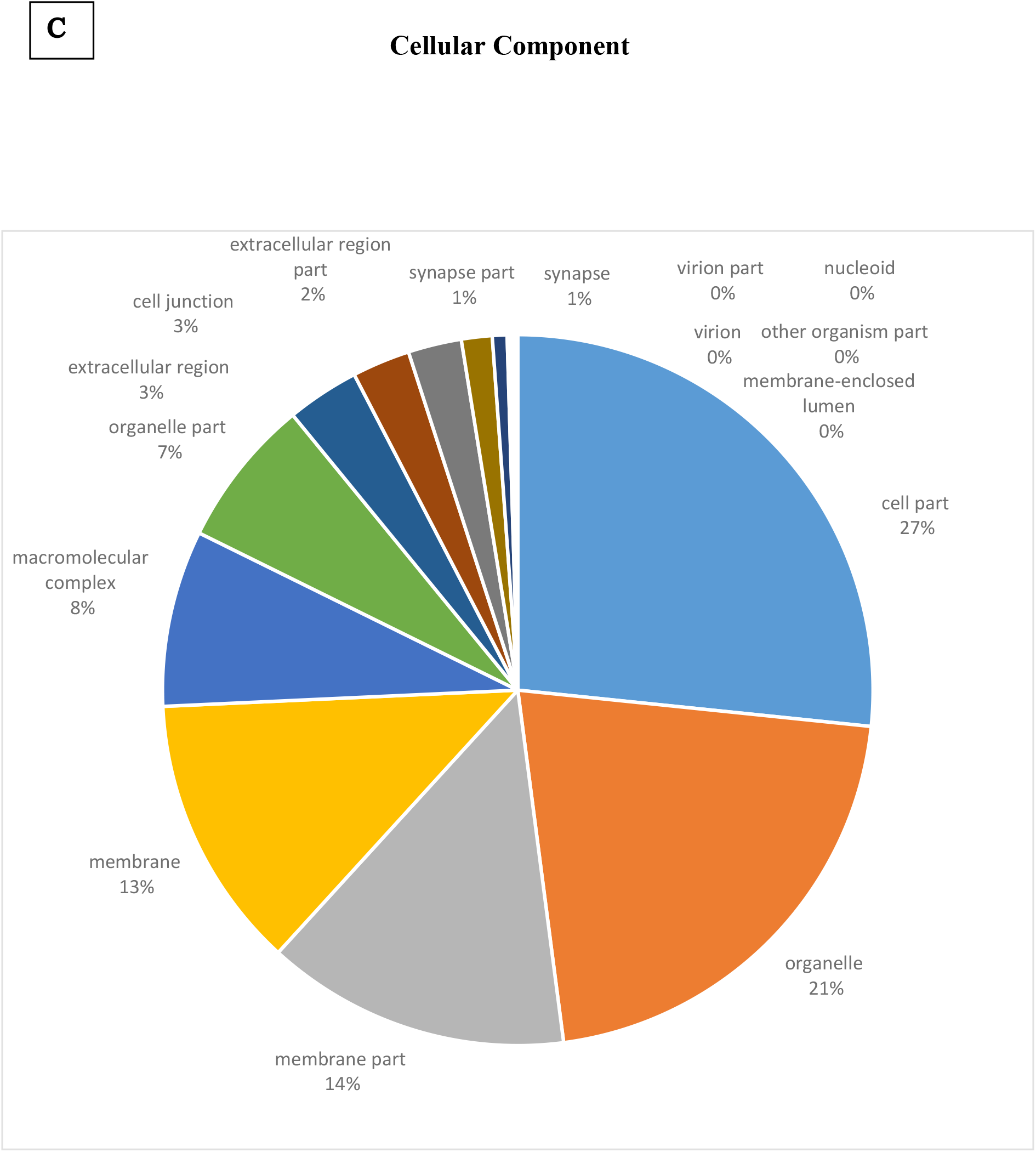
G**O analyses of *Conopomorpha cramerella* transcriptome data.** GO analysis of *Conopomorpha* sequences corresponding to a total of 285,882 contigs that are predicted to be involved in the biological processes (A) and molecular functions (B) and cellular component (C). Classified gene objects are depicted as percentages of the total number of gene objects with GO assignments.

**Figure 5.**
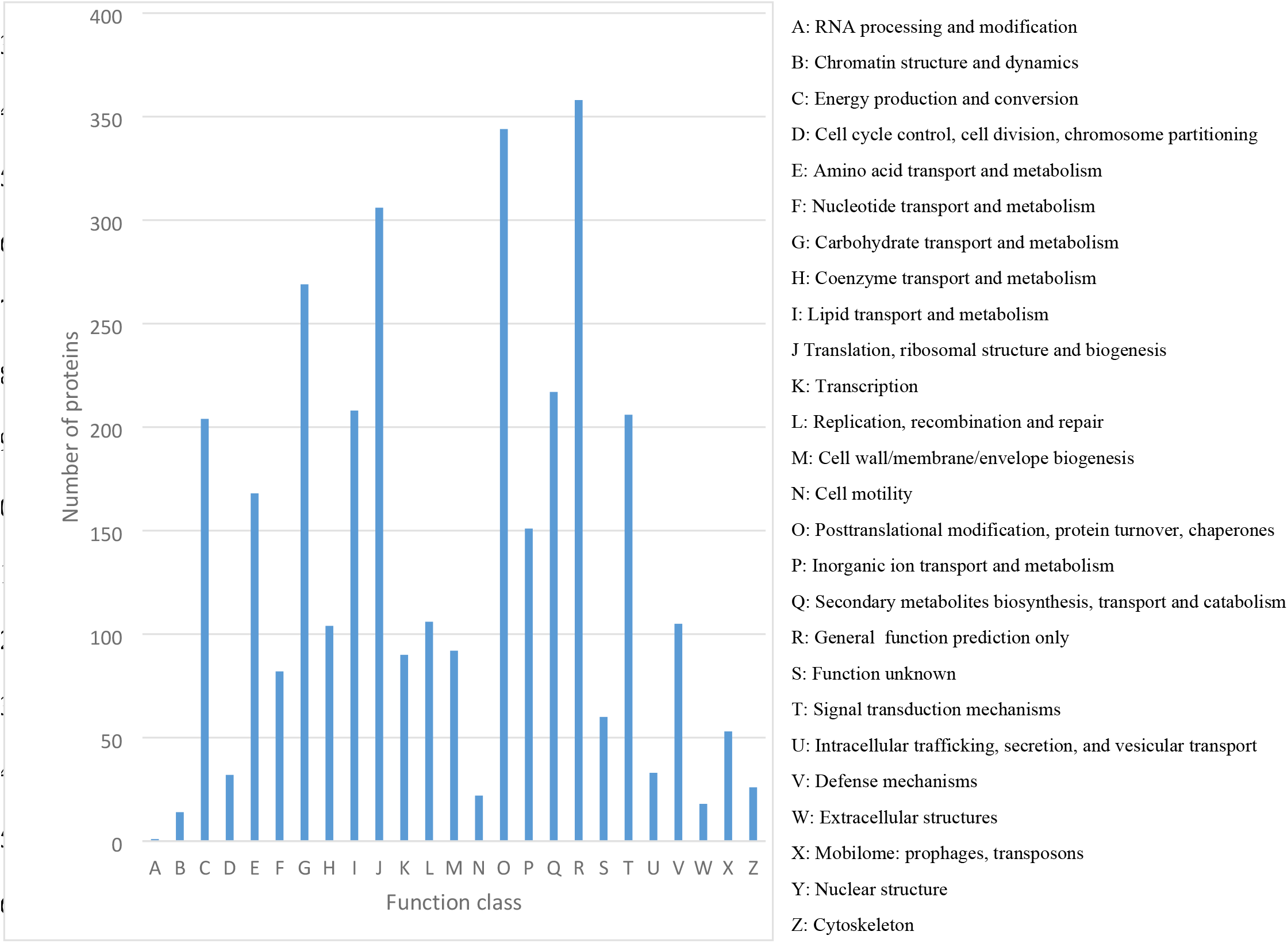
Histogram of clusters of orthologous groups (COG) classification. A total of 3,296 predicted proteins have a COG classification among the 25 categories.

To further evaluate the completeness of our transcriptomic library and the effectiveness of our annotation process, assignments of cluster of orthologous groups (COG) were used. Overall, 3,269 were classified as involved in different metabolic process (Fig. 1). Among the 25 COG categories, the majority of the cluster were “General function prediction only” (358, 10.95%), “Posttranslational modification, protein turnover, chaperones” (344, 10.52%), “Translation, ribosomal structure and biogenesis” (306, 9.36%) and “Carbohydrate transport and metabolism” (296, 8.23%) whereas “RNA processing and modification” (1, 0.03%),

“Chromatin structure and dynamics” (14, 0.43%) and “Extracellular structures” (18, 0.55%) represented the smallest groups (Figure 5).

### Genes involved in general function

Genes involved in general function were listed in Table 2. The results showed that “General function prediction only” constitutes the majority of the cluster within the metabolism pathway classification of the *C. cramerella* transcriptome (Fig. 5). This includes choline dehydrogenase or related flavour protein, GTPase SAR1 family domain, NAD(P)-dependent dehydrogenase, short-chain alcohol dehydrogenase family, pimeloyl-ACP methyl ester carboxylesterase, short-chain dehydrogenase, tetratricopeptide (TPR) repeat and WD40 repeat (Table 2).

**Table 2.** Genes involved in general function

| COG annotation | No. of genes |
| --- | --- |
| Choline dehydrogenase or related flavoprotein | 21 |
| GTPase SAR1 family domain | 49 |
| NAD(P)-dependent dehydrogenase, short-chain alcohol dehydrogenase family | 60 |
| Pimeloyl-ACP methyl ester carboxylesterase | 17 |
| Short-chain dehydrogenase | 13 |
| Tetratricopeptide (TPR) repeat | 11 |
| WD40 repeat | 27 |

Among the genes involved in general function, NAD(P)-dependent dehydrogenase, short- chain alcohol dehydrogenase family has the largest number of genes. Alcohol dehydrogenase is considered a very important enzyme in insect metabolism because it is involved in the catalysis of the reversible conversion of various alcohols in larval feeding sites to their corresponding aldehydes and ketones, thus contributing to detoxification and metabolic purposes [36]. In *Helicoverpa armigera*, alcohol dehydrogenase gene (HaADH5) regulates the expression of CYP6B6, a gene involved in molting and metamorphosis [37]. The second largest group of genes in general function are GTPase. These genes are involved in metabolic pathways of insect [38]. In *Drasophila*, GTPase is found to be involved in endocytosis and vesicle trafficking in the insect renal system [39]. GTPase is also known to regulate diverse cellular and developmental events, by regulating the exocytotic and transcytotic events inside the cell [40]. The third largest group of general function genes are WD40 repeat genes. WD40 proteins are scaffolding molecules in protein-protein interactions and play crucial roles in fundamental biological processes such as the metabolic activities of the insect [41, 42].

### Genes expression profile among the different developmental stages

To identify genes showing differential expression during development, the differentially expressed sequences between two samples were identified (Fig. 6). There were 2,843 differentially expressed genes detected between the larva and pupa samples, including 1,979 down-regulated genes (P<0.05) and 864 up-regulated genes (P<0.05). The large number of differentially expressed genes between these two samples may be attributed to the important molting and metamorphosis processes during transition from larva to pupa. A cascade of physiological processes occurs during molting and complicated physiological processes takes place during metamorphosis including histolysis of larval tissues, remodelling and formation of adult tissues, and a molting cascade similar to the larva molt [43]. In addition, a total of 2,861 differentially expressed genes were detected between adult and larva stage, with 1,646 down-regulated genes and 1,215 up-regulated genes (Figure 6). Between the adult and the pupa stage, 897 genes were down-regulated whereas 1,056 genes were up-regulated from a total of 1,953 differentially expressed genes (Fig. 6).

**Figure 6.**
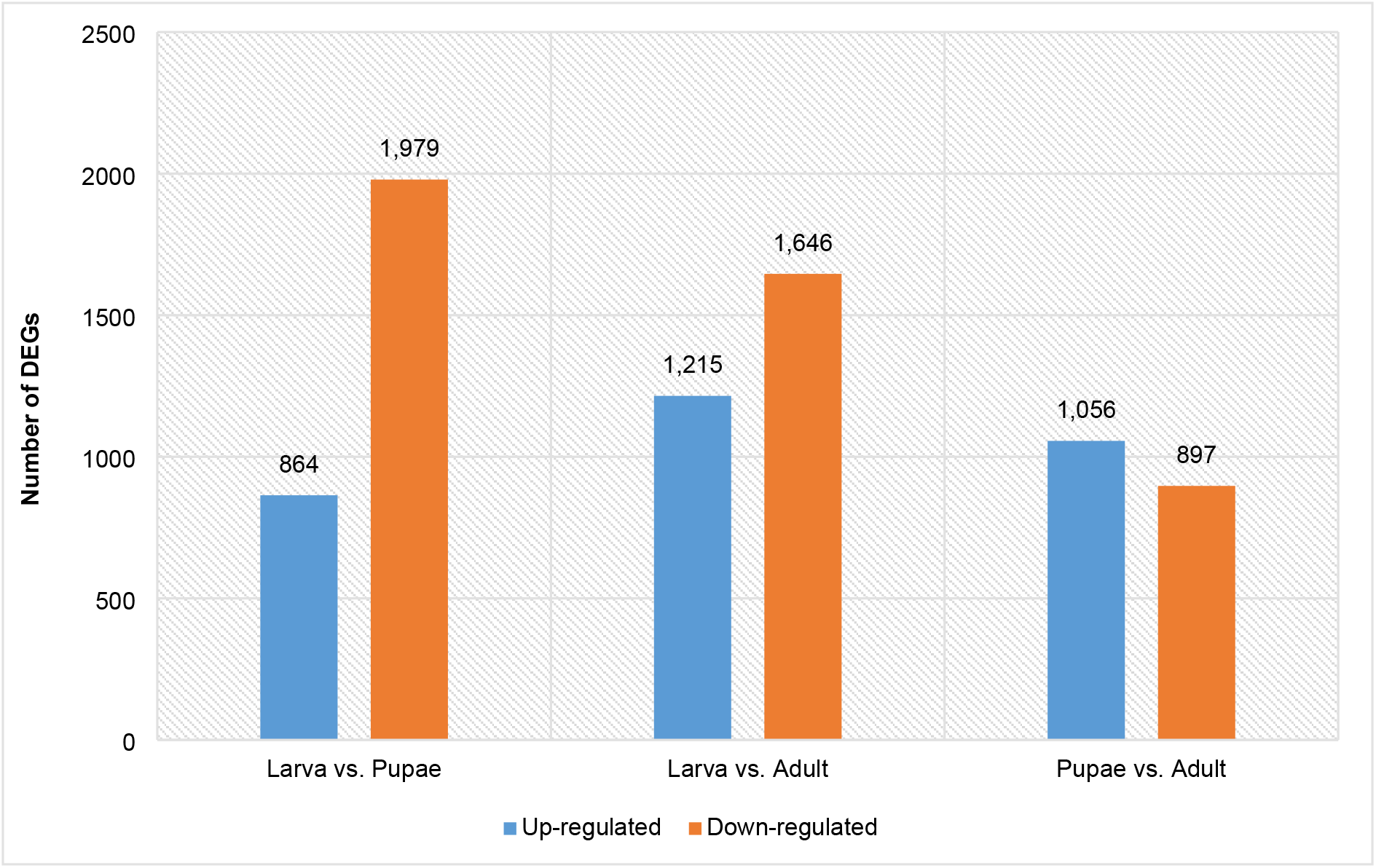
Differentially gene expression profile at different developmental stages. The number of up-regulated and down-regulated genes between larvae and pupae, between adults and pupae, and between adults and larvae are summarized here.

In larva, there is a total of 140,427 expressed genes (>1.0 fpkm), of which 14,023 were known genes and 126,404 novel genes (Table S2). In pupae, a total of 124,368 expressed genes with 13,417 known genes and 110,951 novel genes. In adult moth, the total of expressed genes were 129,652, of which 13,536 were known genes whereas 116,116 were novel genes. The sheer number of novel genes as compared to known genes goes to show that there are many genes in *C. cramerella* that was yet to be discovered.

### Genes involved in reproduction

In insects, sexual reproduction is a very important physiological process and is critical to the maintenance of a population. Therefore, identification of genes involved in reproduction is important and would be helpful for pest control purposes. In addition, it will also be useful to evaluate molecular mechanism for higher order insect’s species. Several reproductive-related genes have been identified (Table 3) in the transcriptome libraries. Among them is the *porin* gene, a male-biased pheromone binding protein, a short chain dehydrogenase/reductase, and a member of the *takeout* gene family [44]. Another reproductive-related genes is the boule (*bol*) gene. This gene is a member of the *Deleted* in *Azoospermia* (DAZ) gene family and plays an important role in meiosis (reductional maturation divisions) in a spermatogenesis of insect male [45]. The gene, *dsx* is also found in the transcriptome analysis. This gene is involved in sex determination in insect [<u>46</u>; <u>47</u>].

**Table 3.** Cocoa pod borer assembled sequences with best-hit matches to insect genes involved in reproductive behaviors

| Gene ID | Insect Gene | Length (bp) | E-value | Protein identity (%) | Function |
| --- | --- | --- | --- | --- | --- |
| TBIU002860 | *porin* | 273 | 3.00E-09 | 67.86 | pheromone binding protein |
| TBIU040283 | *bol* | 1456 | 3.00E-07 | 40.68 | spermatogenesis of insect male |
| TBIU000835 | *dsx* | 344 | 4.00E-09 | 29.91 | sex determination |
| TBIU000002 | *fruitless* | 663 | 1.00E-84 | 92.7 | sex determination |

Another sex-determination gene that is found in *C. cramerella* is the *fruitless* gene. In *Drosophila melanogaster*, the *fruitles*s gene produces sex-specific gene products under the control of the sex-specific splicing cascade and contributes to the formation of the sexually dimorphic circuits [48, 49].

### Verification of differentially expressed genes

In order to evaluate our DEG library, the expression level of seventeen genes involved in development were analysed by qRT-PCR. Results showed that real-time PCR revealed the same expression trend as in the DEG data, albeit with some quantitative differences in expression level (Table 4, Fig. 7). The genes *atr* and *me31B* were highly expressed in the pupa stage. These genes are involved in cross-over patterning effect and transitioning [50, 51] in *Drosophila*, probably plays a crucial role in growth development from larva to pupa. Src64B protein are actively involved in modulating actin level in cell development [52, 53] are highly expressed in the larva. Setdb1 is involved in histone modifications and genome regulation [54] and is expressed higher in the moth compared to the larva stage (Fig. 7). The *pol* gene is almost entirely expressed in the larva and not in the moth. It has function in RNA synthesis and as a growth effector of Ras/ERK signalling in *Drosophila* [55]. As control, actin is used as it is demonstrated to be almost equally expressed in all the three developmental stages (Fig. 7). These data will provide us with molecular targets to further study on the development of *Conopomorpha cramerella*.

**Figure 7.**
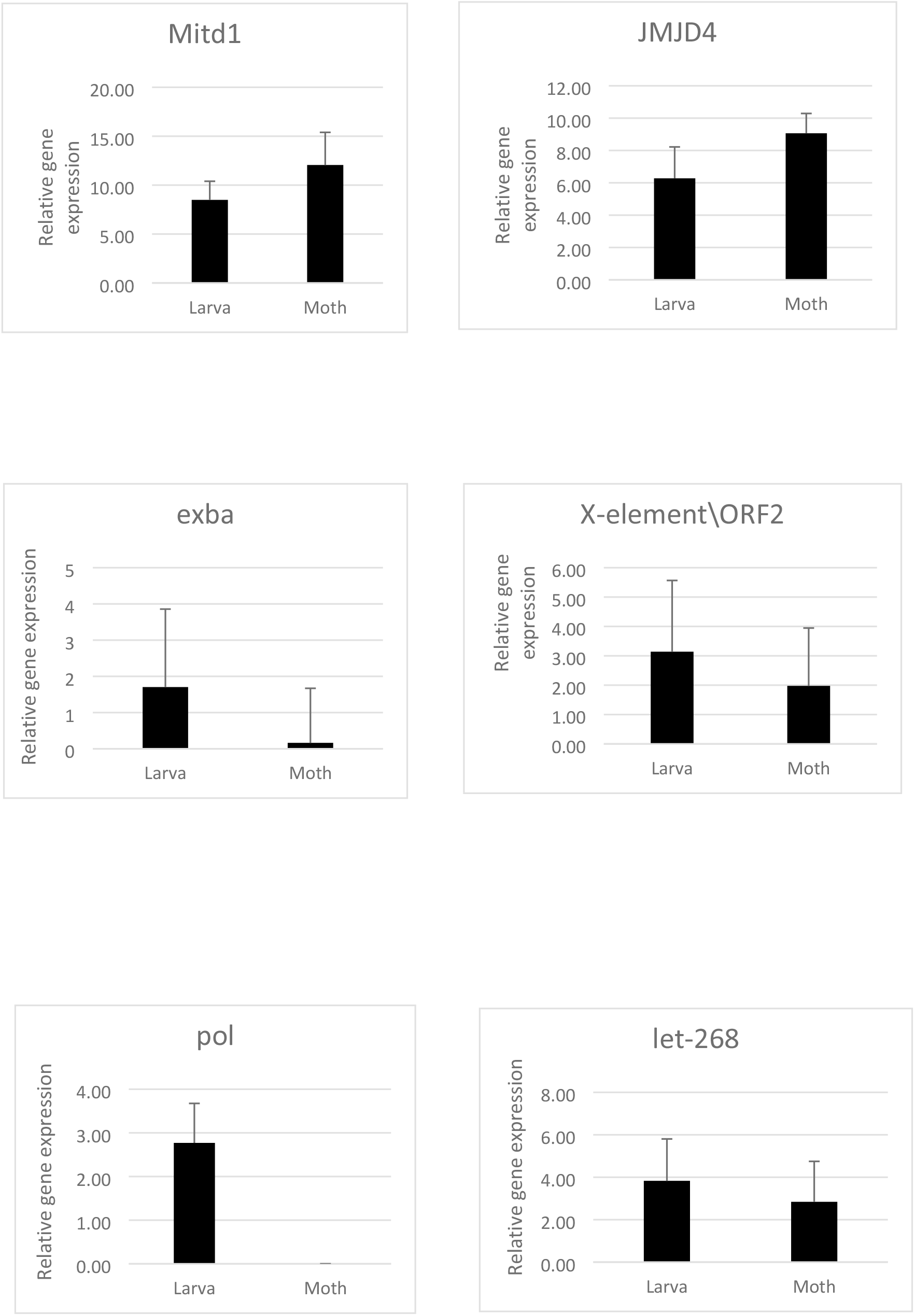
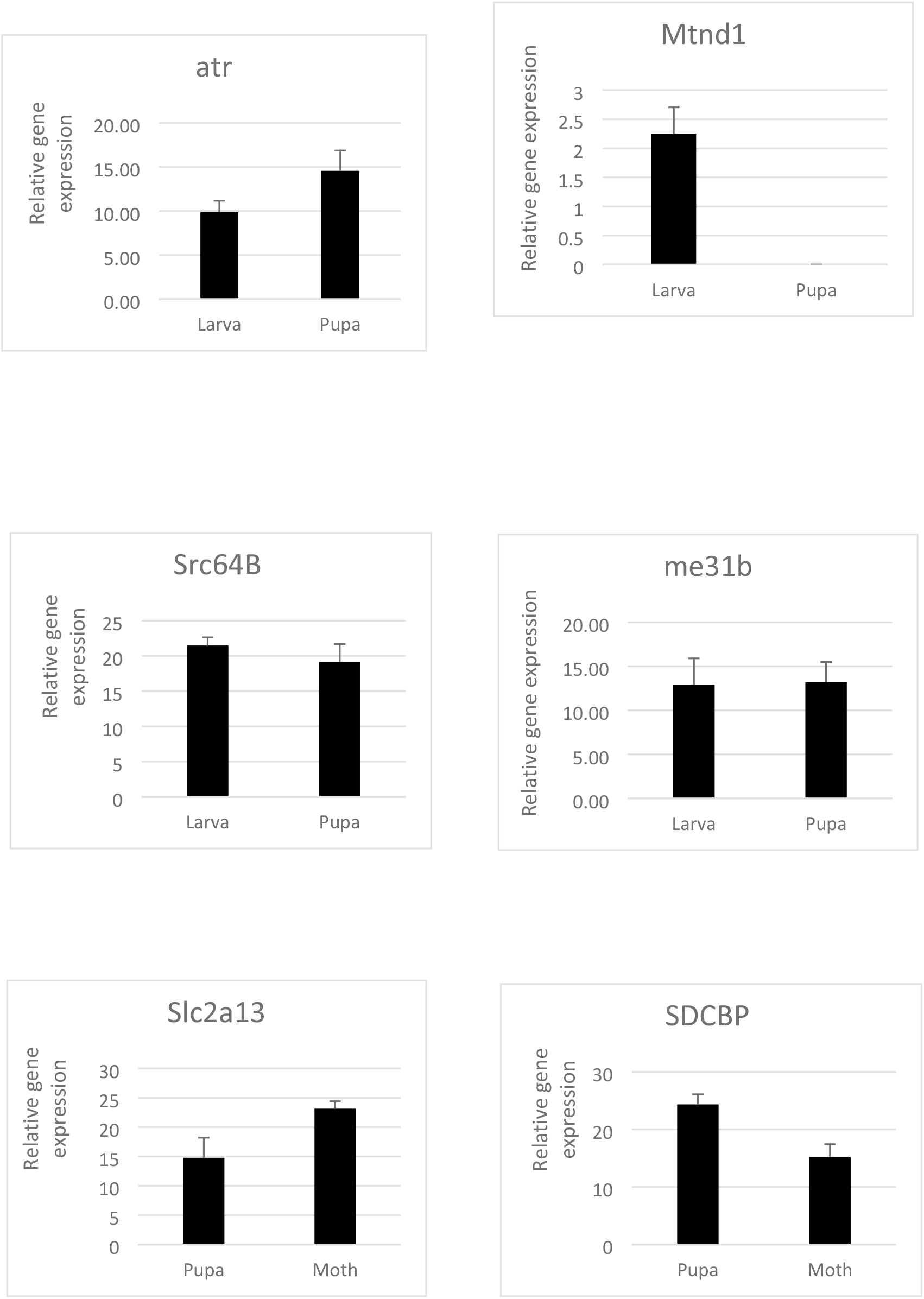
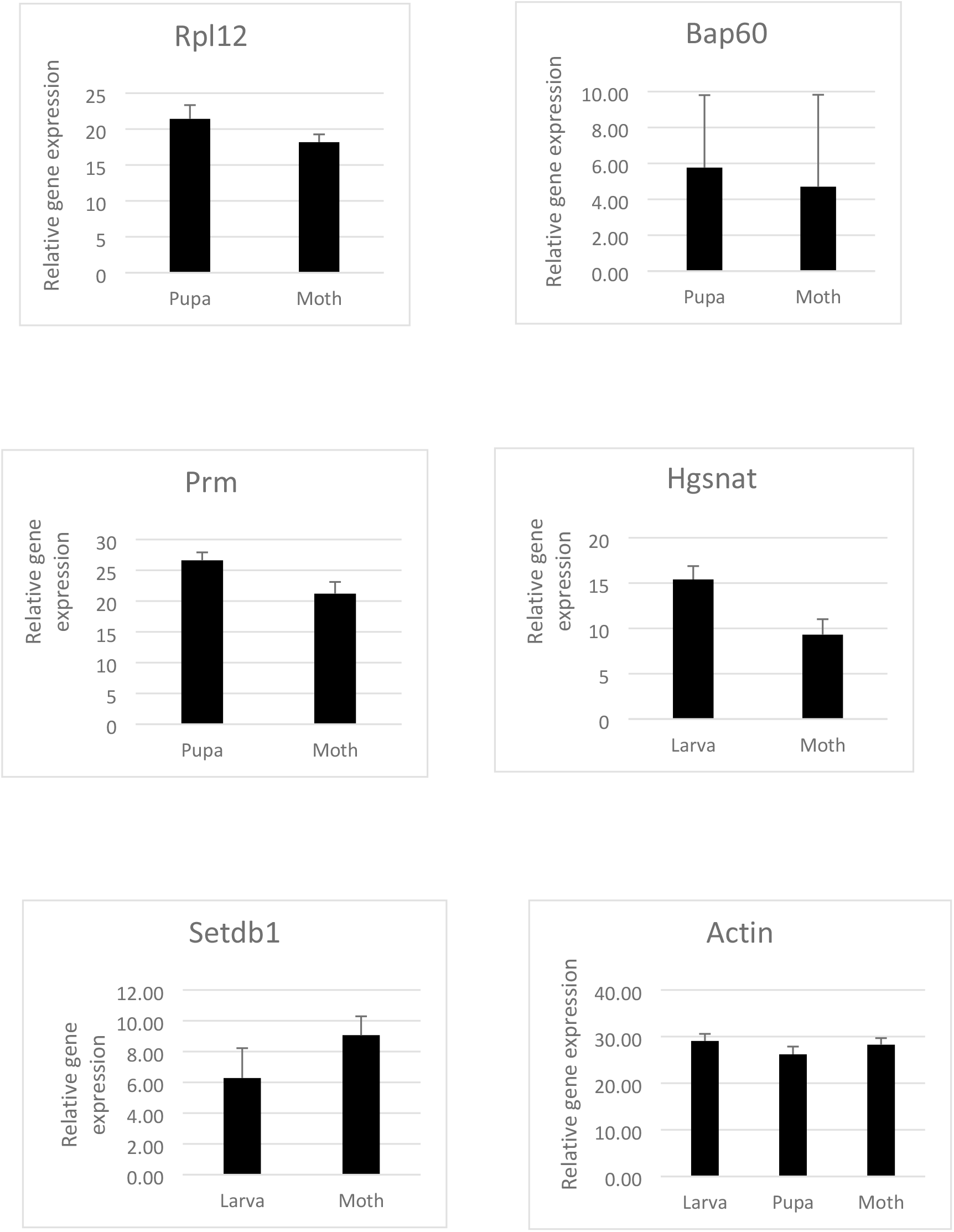
Q**R**T**-PCR validation of the differentially expressed genes between each of the two stages of growth (larva vs. pupa, pupa vs. moth and larva vs. moth).** Relative transcript levels are calculated by real-time PCR using Actin gene as reference standard. Three biological replicates were performed, and the data shown are typical results.

**Table 4.**
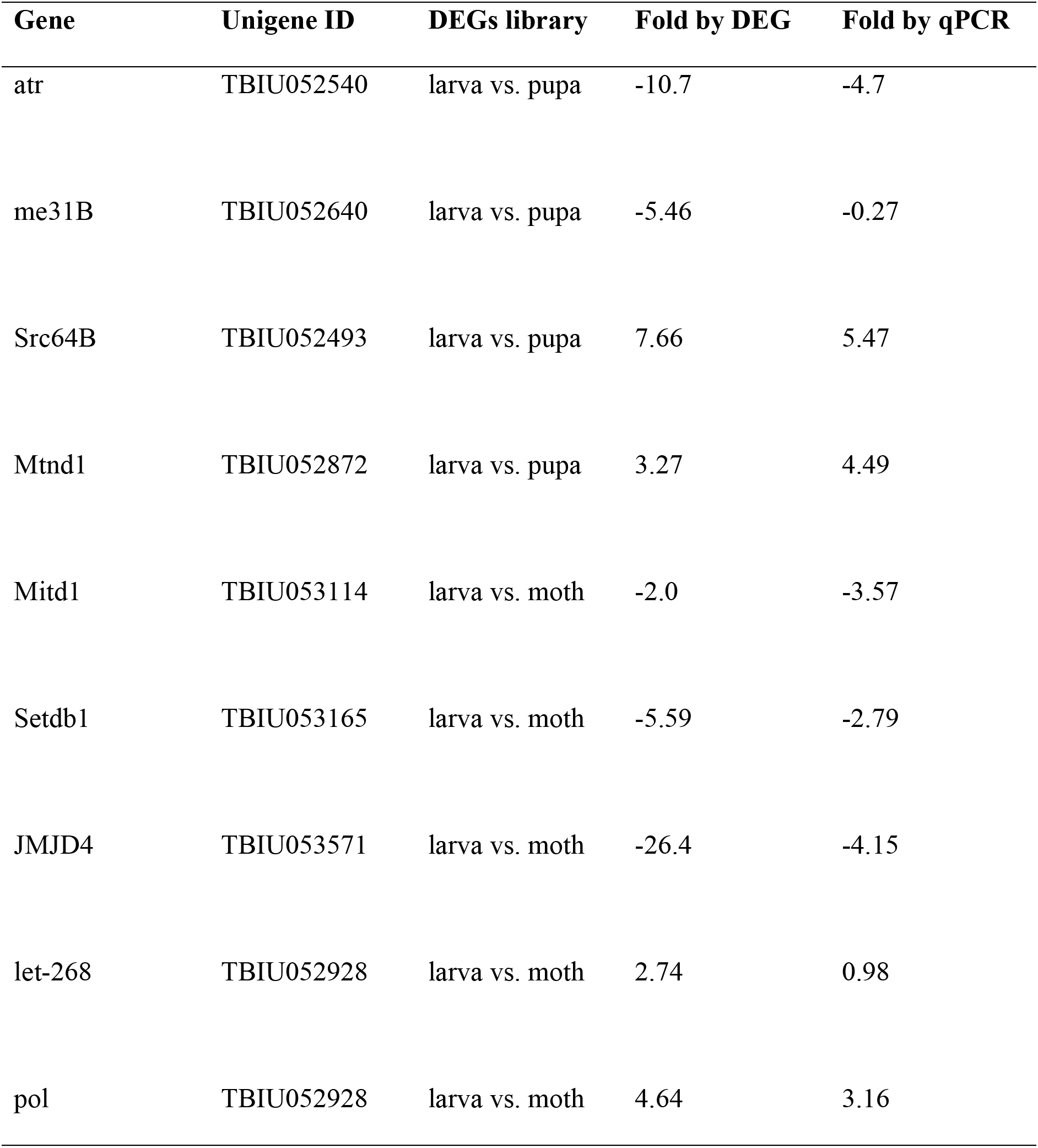
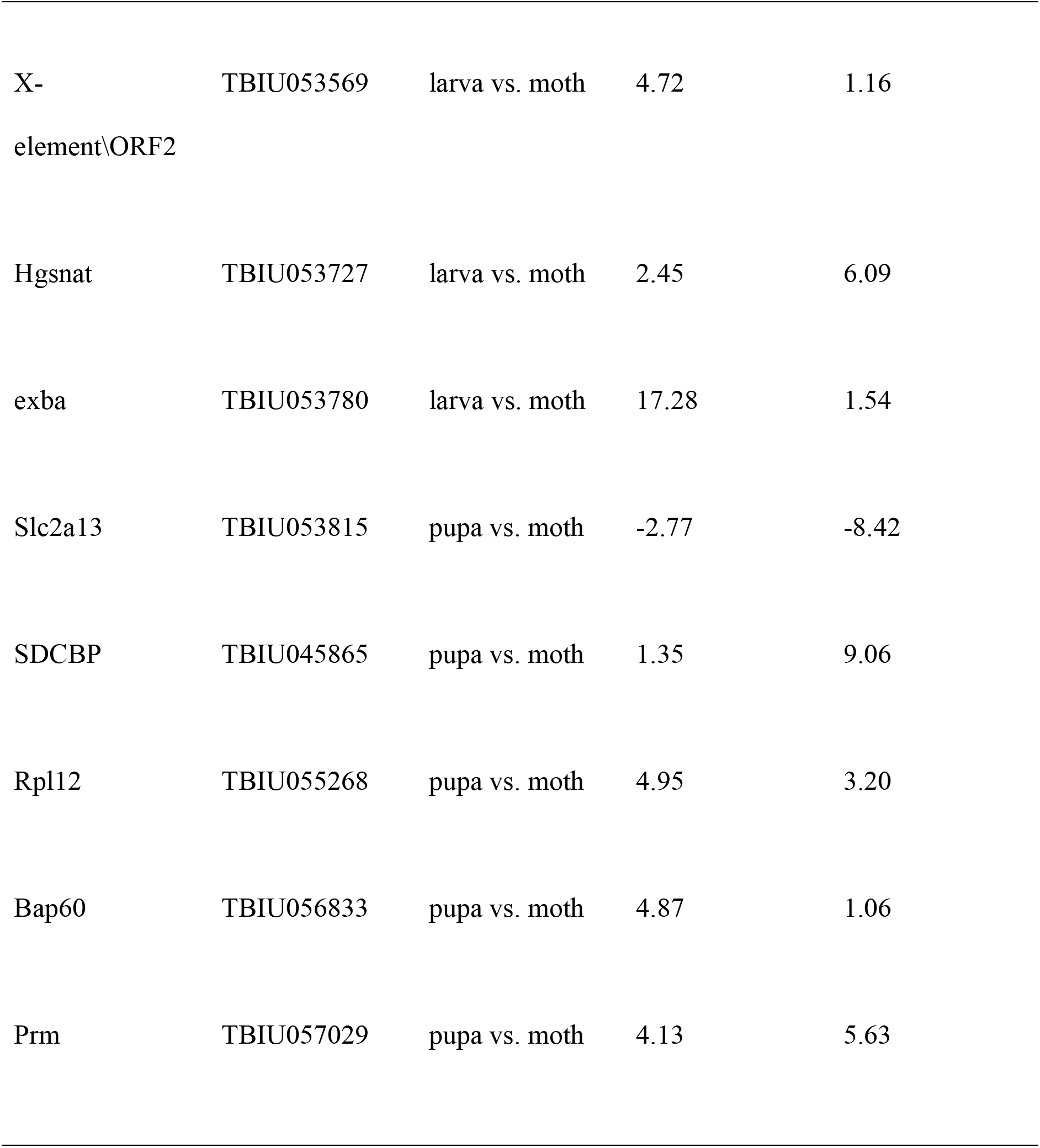
Comparisons of DEGs data and qRT-PCR results

| Gene | Unigene ID | DEGs library | Fold by DEG | Fold by qPCR |
| --- | --- | --- | --- | --- |
| atr | TBIU052540 | larva vs. pupa | -10.7 | -4.7 |
| me31B | TBIU052640 | larva vs. pupa | -5.46 | -0.27 |
| Src64B | TBIU052493 | larva vs. pupa | 7.66 | 5.47 |
| Mtnd1 | TBIU052872 | larva vs. pupa | 3.27 | 4.49 |
| Mitd1 | TBIU053114 | larva vs. moth | -2.0 | -3.57 |
| Setdb1 | TBIU053165 | larva vs. moth | -5.59 | -2.79 |
| JMJD4 | TBIU053571 | larva vs. moth | -26.4 | -4.15 |
| let-268 | TBIU052928 | larva vs. moth | 2.74 | 0.98 |
| pol | TBIU052928 | larva vs. moth | 4.64 | 3.16 |
| X-element\ORF2 | TBIU053569 | larva vs. moth | 4.72 | 1.16 |
| Hgsnat | TBIU053727 | larva vs. moth | 2.45 | 6.09 |
| exba | TBIU053780 | larva vs. moth | 17.28 | 1.54 |
| Slc2a13 | TBIU053815 | pupa vs. moth | -2.77 | -8.42 |
| SDCBP | TBIU045865 | pupa vs. moth | 1.35 | 9.06 |
| Rpl12 | TBIU055268 | pupa vs. moth | 4.95 | 3.20 |
| Bap60 | TBIU056833 | pupa vs. moth | 4.87 | 1.06 |
| Prm | TBIU057029 | pupa vs. moth | 4.13 | 5.63 |

## Conclusions

We have generated a comprehensive transcriptome of the *C. cramerella* development using Illumina NovaSeq6000 platform. The single run produced 285,882 contigs with a mean length of 374 bp. A large number of genes involved in reproduction, general function and development pathways are found in the transcriptome. In addition, genes differentially expressed at different development stages were identified. To our knowledge, this is the first report of transcriptome sequencing in *C. cramerella*, a lepidopteran insect pest lacking a reference genome. These data make a substantial contribution to genetic resources of cocoa pod borer. It also provide potential molecular targets for the control of *C. cramerella* using RNAi. Finally, the study may also aid in the understanding of the molecular basis of development and reproduction in cocoa pod borer insect.

## Acknowledgement

We would like to thank the Director-General of the Malaysia Cocoa Board for permission to publish this paper. We also like to thank the Director of Biotechnology for allowing funding from the 11th Malaysia Development Fund for this project. Lastly, we also thank Neoscience Sdn. Bhd., Malaysia and Theragen, South Korea for the sequencing work.

## Author Contributions

Conceived and designed the experiment: CLT, RK, WWL, WML. Performed the experiment: CLT, WML. Analysed the data: CLT, WWL, WML. Wrote the paper: CLT, WML.

## Supporting information

**Figure S1.**
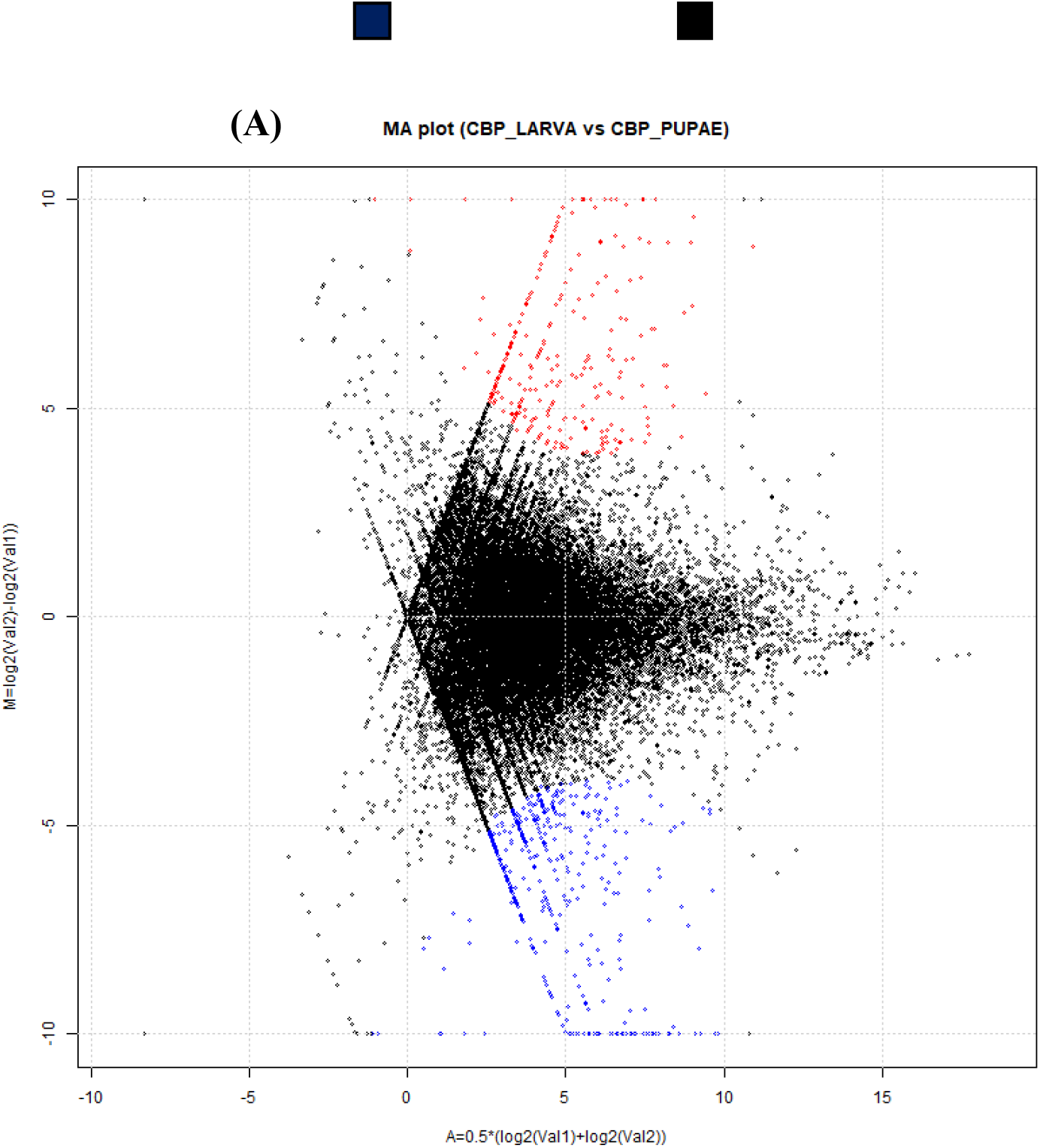
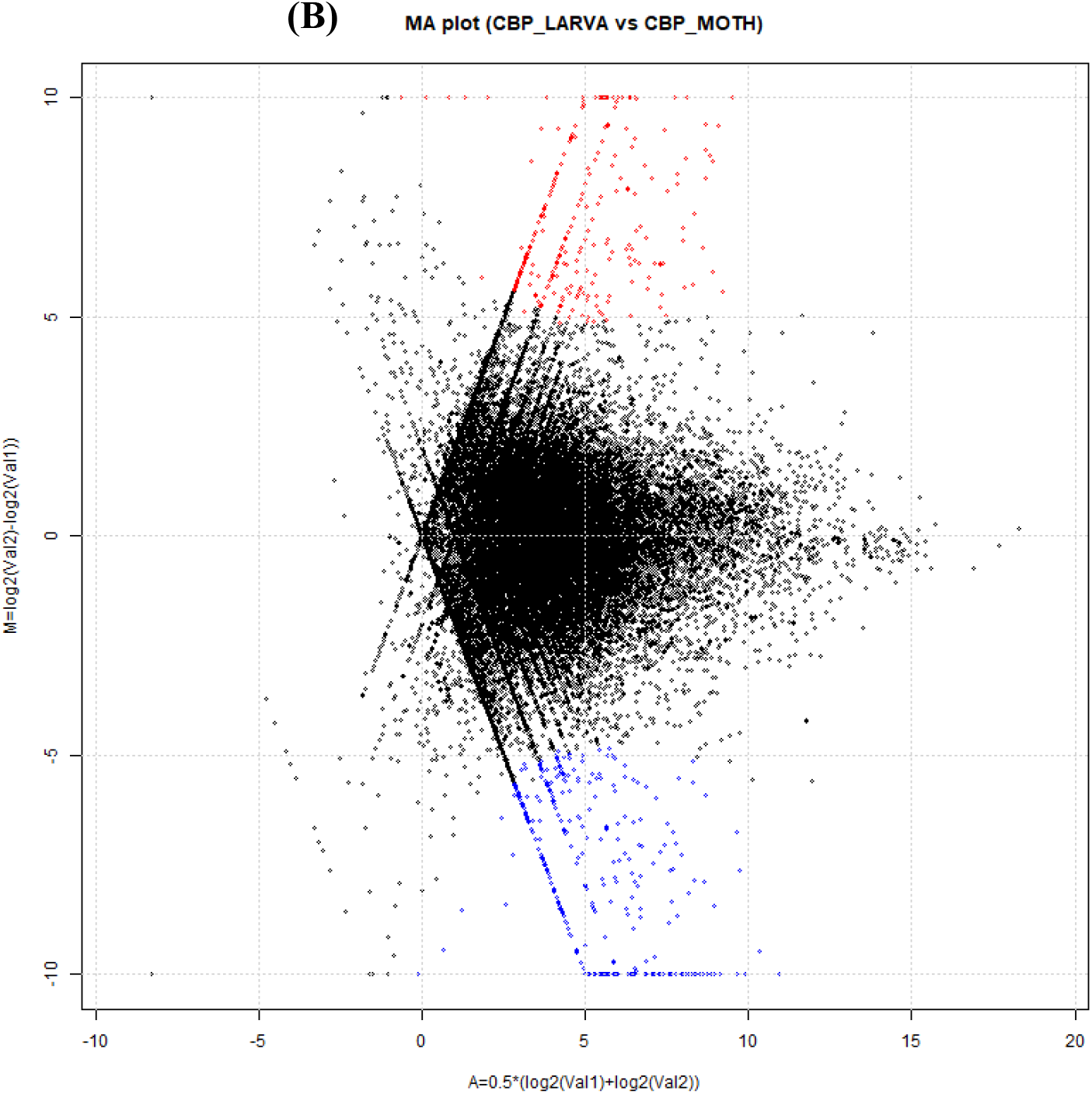
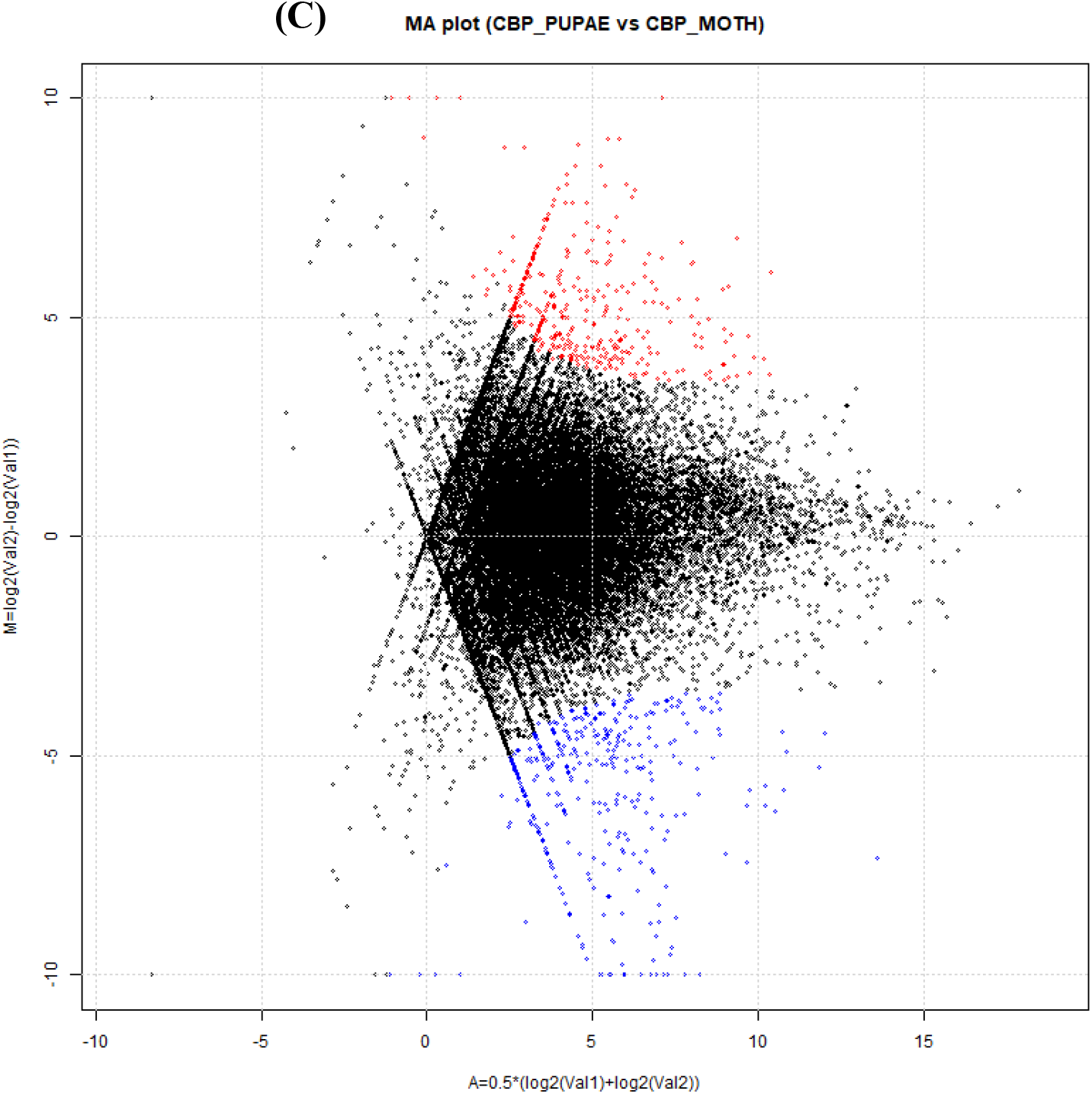
Comparison of sequence expression between the larvae and pupae (A), moth and larvae (B), as well as moth and pupae (C). The abundance of each gene was normalised as Fragments Per Kilobase per Million (FPKM). The differentially expressed genes are shown in red and blue, while the other genes that are not differentially expressed (not DEGs) are 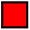 Up-regulated Down regulated Not DEGs shown in black.

**Table S1.**
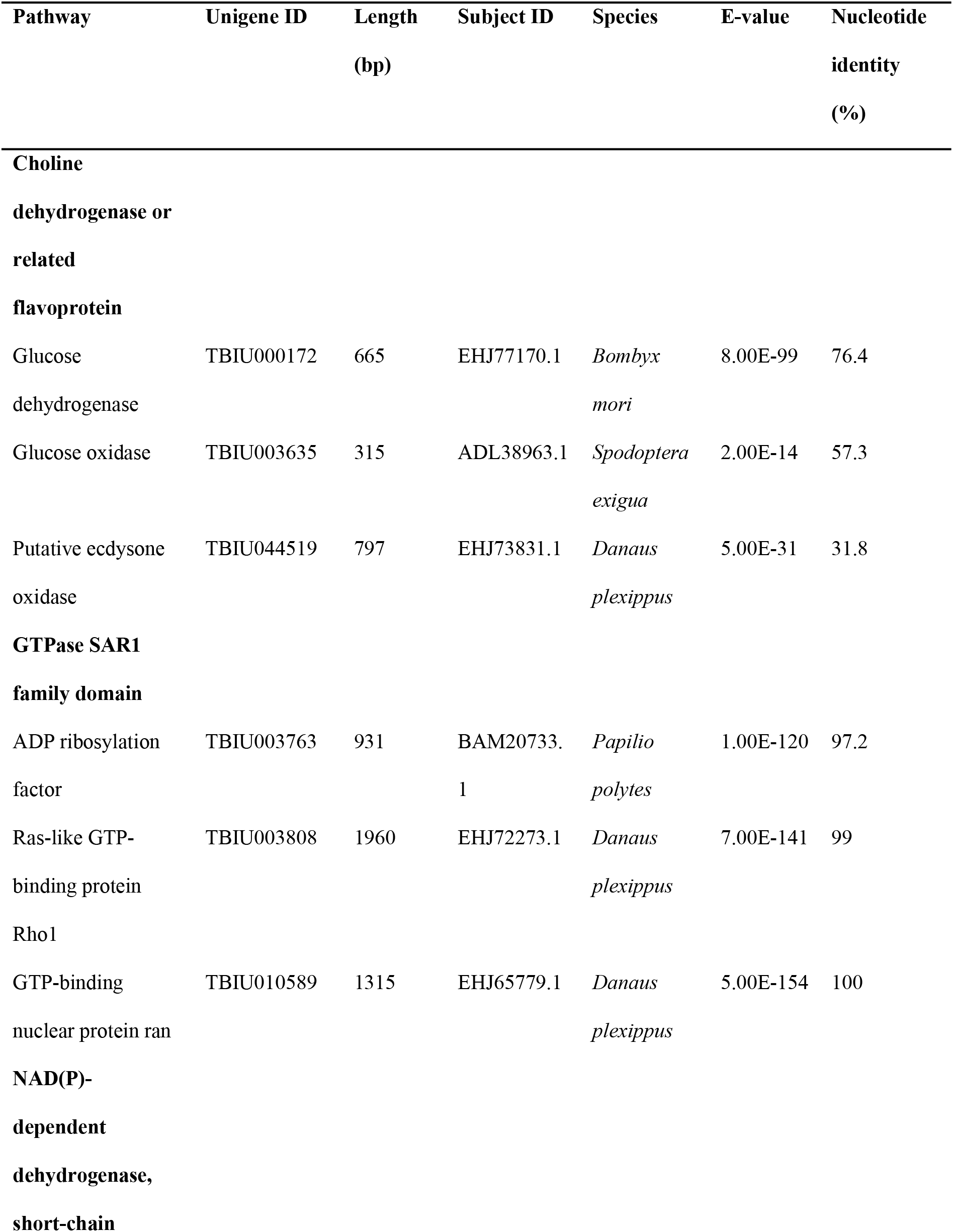
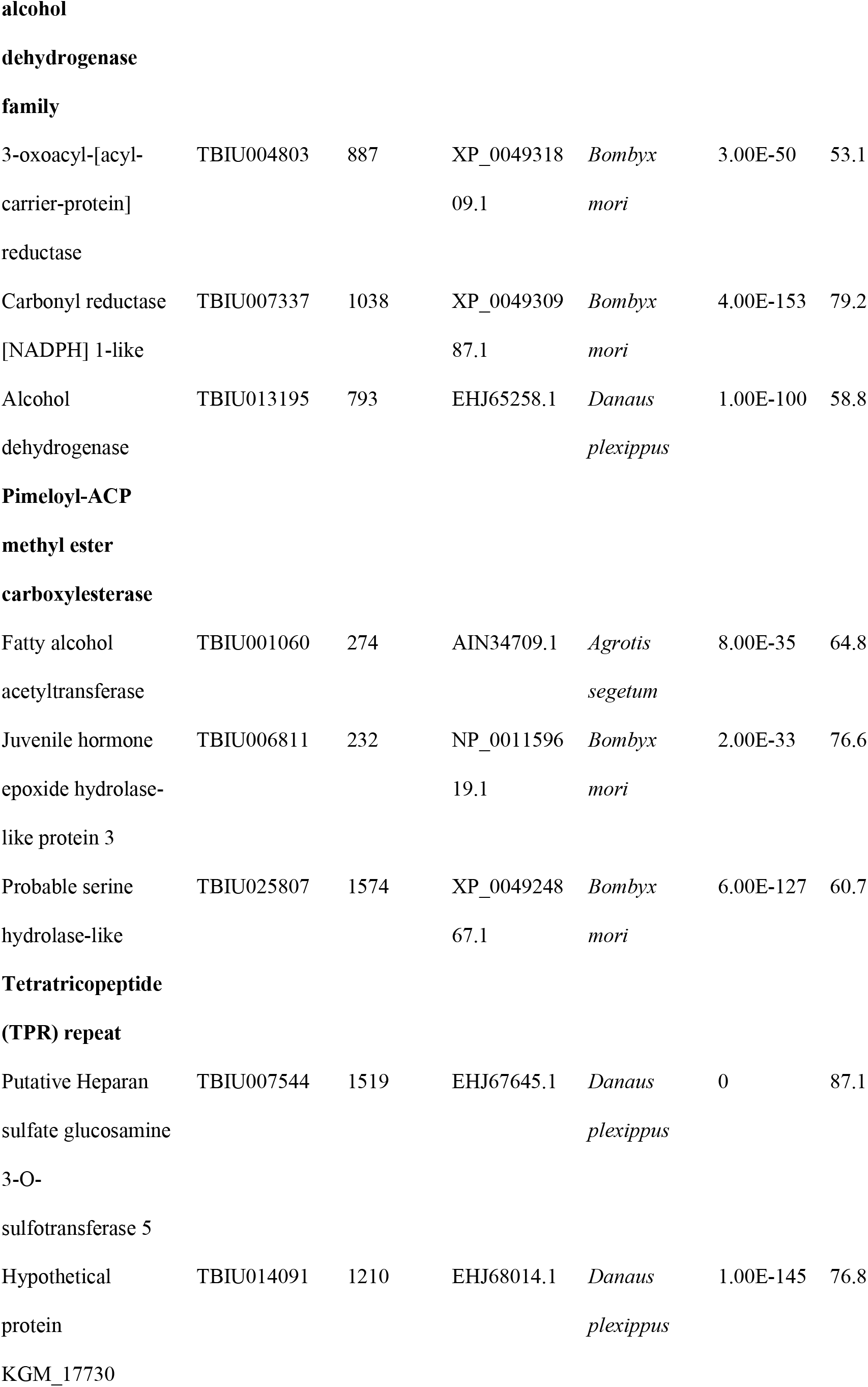
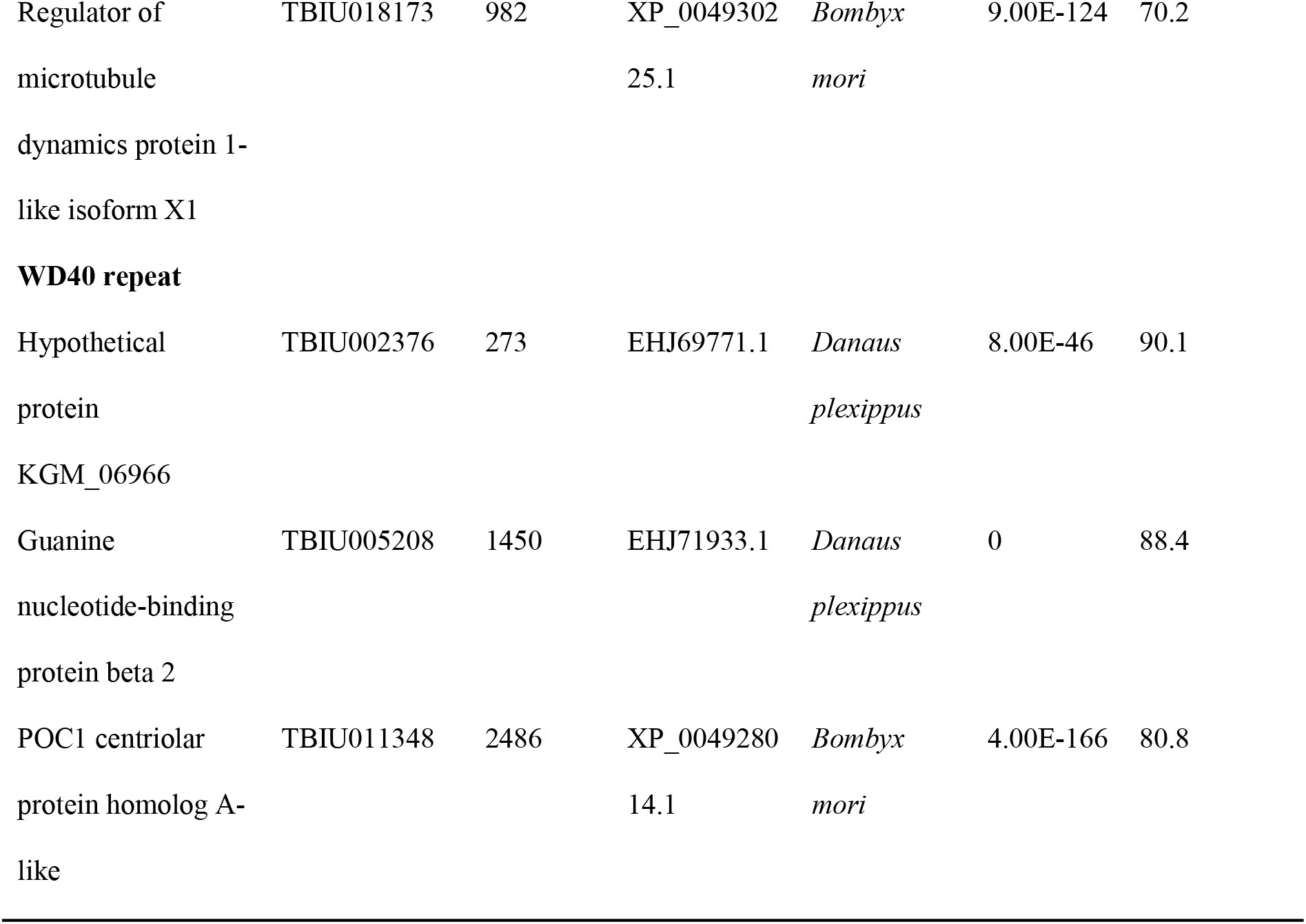
Selected general function genes identified in the *Conopomorpha cramerella* transcriptome with best-hit matches to other insects

**Table S2.** Expression of genes in three different development stages of *Conopomorpha cramerella*.

| Name | Gene |  |  |  | Gene (> FPKM 1.0) |  |  |  |
| --- | --- | --- | --- | --- | --- | --- | --- | --- |
|  | Expressed | Known | Novel | Unexpressed | Expressed | Known | Novel | Unexpressed |
| Larva | 143,065 | 14,236 | 128,829 | 142,817 | 140,427 | 14,023 | 126,404 | 74,966 |
| Pupae | 125,651 | 13,518 | 112,133 | 160,231 | 124,368 | 13,417 | 110,951 | 91,025 |
| Moth | 131,120 | 13,689 | 117,431 | 154,762 | 129,652 | 13,536 | 116,116 | 85,741 |

**Table S3.** Selected up-regulated and down regulated genes between samples (larva vs. pupa).

| Gene name | Unigene ID | Gene description | Source | Expression<br>– Larva<br>(FPKM) | Expression<br>– Pupa<br>(FPKM) |
| --- | --- | --- | --- | --- | --- |
| Up-regulated |  |  |  |  |  |
| genes |  |  |  |  |  |
| atr | TBIU052540 | Serine/threonine-<br>protein kinase atr | SWISS;ACC :Q9DE14 | 0 | 10.7 |
| me31B | TBIU052640 | Putative ATP-dependent RNA<br>helicase me31b | SWISS;ACC :P23128 | 0 | 5.46 |
| dtwd1 | TBIU052700 | DTW domain-containing protein 1 | SWISS;ACC :Q6DDV1 | 2.67 | 3.52 |
| Trappc2 | TBIU052984 | Trafficking protein<br>particle complex<br>subunit 2 | SWISS;ACC :D3ZVF4 | 0 | 1.64 |
| AGBL1 | TBIU052987 | Cytosolic<br>carboxypeptidase 4 | SWISS;ACC :Q96MI9 | 0 | 5.25 |
| Down-regulated |  |  |  |  |  |
| genes |  |  |  |  |  |
| Ubr5 | TBIU052459 | E3 ubiquitin-protein<br>ligase | SWISS;ACC :Q62671 | 14.93 | 6.65 |
| Src64B | TBIU052493 | Tyrosine-protein kinase<br>Src64B | SWISS;ACC :P00528 | 7.66 | 0 |
| Dnah2 | TBIU052596 | Dynein heavy chain 2,<br>axonemal | SWISS;ACC :P0C6F1 | 2.16 | 0 |
| pycr1 | TBIU046555 | Pyrroline-5-carboxylate<br>reductase 3 | SWISS;ACC<br>:Q5SPD7 | 5.18 | 2.49 |
| Mtnd1 | TBIU052872 | NADH-ubiquinone<br>oxidoreductase chain 1 | SWISS;ACC<br>:P03888 | 3.27 | 0 |

**Table S4.** Selected up-regulated and down regulated genes between samples (larva vs. moth).

| Gene name | Unigene ID | Gene description | Source | Expression<br>– Larva<br>(FPKM) | Expression<br>– Moth<br>(FPKM) |
| --- | --- | --- | --- | --- | --- |
| **Up-regulated** |  |  |  |  |  |
| **genes** |  |  |  |  |  |
| Mitd1 | TBIU053114 | MIT domain-containing protein 1 | SWISS;ACC<br>:Q5I0J5 | 5.28 | 7.28 |
| Setdb1 | TBIU053165 | Histone-lysine N-methyltransferase<br>SETDB1 | SWISS;ACC<br>:O88974 | 0 | 5.59 |
| JMJD4 | TBIU053571 | JmjC domain-containing protein 4 | SWISS;ACC<br>:Q5ZHV5 | 15.12 | 41.52 |
| NDUFS5 | TBIU053671 | NADH dehydrogenase<br>[ubiquinone] iron-<br>sulfur protein 5 | SWISS;ACC<br>:Q0MQH3 | 342.69 | 967.27 |
| RTase | TBIU054001 | Probable RNA-directed<br>DNA polymerase from<br>transposon BS | SWISS;ACC<br>:Q95SX7 | 0 | 2.17 |
| **Down-regulated** |  |  |  |  |  |
| **genes** |  |  |  |  |  |
| let-268 | TBIU052928 | Procollagen-lysine,2-oxoglutarate 5-dioxygenase | SWISS;ACC<br>:Q20679 | 2.74 | 0 |
| pol | TBIU052994 | RNA-directed DNA<br>polymerase from<br>mobile element jockey | SWISS;ACC<br>:P21328 | 4.64 | 0 |
| X-element\ORF2 | TBIU053569 | Probable RNA-directed<br>DNA polymerase from<br>transposon X-element | SWISS;ACC<br>:Q9NBX4 | 5.93 | 1.21 |
| Hgsnat | TBIU053727 | Heparan-alpha-<br>glucosaminide N-acetyltransferase | SWISS;ACC<br>:Q3UDW8 | 4.82 | 2.37 |
| exba | TBIU053780 | Protein extra bases | SWISS;ACC<br>:Q9VNE2 | 17.28 | 0 |

**Table S5.** Selected up-regulated and down regulated genes between samples (pupa vs. moth).

| Gene name | Unigene ID | Gene description | Source | Expression<br>– Pupa<br>(FPKM) | Expression<br>– Moth<br>(FPKM) |
| --- | --- | --- | --- | --- | --- |
| **Up-regulated** |  |  |  |  |  |
| **genes** |  |  |  |  |  |
| RpS25 | TBIU052930 | 40S ribosomal protein<br>S25 | SWISS;ACC<br>:Q962Q5 | 0 | 2.77 |
| Slc2a13 | TBIU053815 | Proton myo-inositol<br>cotransporter | SWISS;ACC<br>:Q3UHK1 | 1.91 | 22 |
| alg9 | TBIU054025 | Alpha-1,2-mannosyltransferase<br>alg9 | SWISS;ACC<br>:Q9P7Q9 | 0 | 2.03 |
| ABCF2 | TBIU054126 | ATP-binding cassette<br>sub-family F member 2 | SWISS;ACC<br>:Q2KJA2 | 0 | 13.25 |
| zc3h15 | TBIU054262 | Zinc finger CCCH<br>domain-containing<br>protein 15 | SWISS;ACC<br>:Q803J8 | 0 | 2.21 |
| **Down-regulated** |  |  |  |  |  |
| **genes** |  |  |  |  |  |
| SDCBP | TBIU045865 | Syntenin-1 | SWISS;ACC<br>:O00560 | 1.35 | 0 |
| Rpl12 | TBIU055268 | 60S ribosomal protein<br>L12 | SWISS;ACC<br>:P35979 | 4.95 | 0 |
| 4CLL4 | TBIU056797 | 4-coumarate--CoA<br>ligase-like 4 | SWISS;ACC<br>:P0C5B6 | 6.99 | 0 |
| Bap60 | TBIU056833 | Brahma-associated<br>protein of 60 kDa | SWISS;ACC<br>:Q9VYG2 | 4.87 | 0 |
| Prm | TBIU057029 | Paramyosin, long form | SWISS;ACC<br>:P35415 | 4.13 | 0 |

## Notes

### Competing Interest Statement

The authors have declared no competing interest.

